# High-intensity interval exercise affects explicit sequential motor consolidation with both physical and mental practice

**DOI:** 10.1101/2025.05.10.653061

**Authors:** Guillaume Digonet, Thomas Lapole, Juliette Gelebart, Jérémie Bouvier, Lucie Blanco, Justine Magaud, Vincent Pialoux, Maxime Pingon, Emeric Stauffer, Ursula Debarnot

## Abstract

High-intensity interval exercise (HIIE) is known to enhance motor consolidation following physical practice (PP), but its effects on sequential motor learning (SML) through PP or motor imagery (MI) remain unclear. We examined whether HIIE modulates SML consolidation in 48 participants who learned an explicit SML task through PP or MI. Performance was assessed before and after acquisition, after HIIE or rest, and at 24 hours and 7 days. Both PP and MI improved performance, with greater gains for PP (p = 0.042), and both induced intracortical disinhibition (p = 0.03). HIIE increased BDNF (p = 0.044) and lactate levels (p < 0.001), markers typically linked to neuroplasticity, yet unexpectedly impaired SML at early (p < 0.01) and late consolidation (p < 0.05), without affecting excitability. These findings challenge the presumed coupling between exercise-induced biomarkers and behavioral gains, suggesting that HIIE may hinder consolidation when explicit components of motor learning are involved.

**Significance Statement:** HIIE is increasingly proposed as a tool to boost neuroplasticity and enhance motor learning. However, whether its benefits extend to all forms of learning remains unclear. Here, we show that both physical and motor imagery practice improve SML and induce intracortical disinhibition, a neurophysiological signature of plasticity. Surprisingly, HIIE impaired SML consolidation at both early and late stages, despite increases in BDNF and lactate, biomarkers typically linked to learning facilitation. This deterioration was observed across both practice modalities and is likely driven by the explicit, cognitively demanding nature of the task. These findings challenge the generalizability of HIIE’s beneficial effects and highlight the need to align exercise-based interventions with the specific cognitive-motor demands of the learning task. Such insights are critical for optimizing motor learning strategies in both athletic training and neurorehabilitation.

## Main Text

### Introduction

Sequential motor skill learning (SML), such as a finger tapping task, is typically characterized by an online acquisition phase requiring repetitive task practice, followed by an offline consolidation phase wherein motor memory traces are reorganized into stabilized or even enhanced representations without further practice^1,2^. Acquisition of a SML task is commonly realized by repetitive physical practice (PP), however, it can also be accomplished with motor imagery practice (MI), i.e. mental state during which the cognitive representation of motor movement is rehearsed without engaging its physical execution^3^. Accumulated evidence demonstrated that MI is a cost-effective and easily feasible substitute for executed movement as a means to activate motor networks, making it an interesting alternative to PP, especially in sport and motor (re)learning contexts^4^. Further evidence suggested that SML can be consolidated through both PP and MI from early to longer timescale periods i.e., within minutes to week^5,6^. From a neurophysiological perspective, motor skill acquisition with PP has been associated to neural plasticity as evidenced by increased amplitude of motor evoked potential (MEPs) elicited by transcranial magnetic stimulation (TMS)^7,8^. Yet such finding appears equivocal, as some studies reported either reductions or no changes in MEPs amplitude^9-11^. A more consistent result is the reduction in short intracortical inhibition (SICI) right after PP acquisition, as measured through a paired-pulse paradigm^11-13^. When considering acquisition with MI, the literature conversely reported no change in MEPs nor SICI following motor skill learning through mental practice^14-16^. To date, however, the behavioral and neural plasticity associated with PP or MI during SML acquisition and consolidation remains largely underexplored^17^.

Recent evidence suggested that performing a single session of high-intensity interval exercise (HIIE), that alternates short periods of intense exercise with active recovery, may be a powerful approach for enhancing motor skill consolidation and promoting neuroplasticity^18^. While these benefits are well-documented for visuomotor tasks^19^, few studies have examined its effect on SML paradigms that depend more heavily on explicit memory components^20-22^. Findings from these behavioral studies did not reveal any additional gains in performance following HIIE, hence suggesting that physical exercise might not contribute to further improving consolidation of explicit SML acquired with PP. Conversely, Monany et al.^23^ demonstrated enhanced consolidation of an explicit SML task acquired with MI when followed by a moderate-intensity exercise. These contrasting findings highlight the need to further examine whether physical exercise may influence consolidation of explicit SML acquired with PP and MI.

The theoretical rationale behind the effect of HIIE suggests that elevated lactate production stimulates BDNF release, with both molecules contributing to neuroplasticity and memory consolidation^24-26^. However, experimental evidence linking lactate and BDNF to enhanced motor consolidation remains limited, with only one study reporting positive correlations between their release and improvements in both early- and late-consolidation of visuomotor tracking performance^27^. Notably, lactate elevation induced by high-intensity exercise has been associated with increased corticospinal excitability^28^, suggesting a potential mechanism through which high-intensity may influence motor learning. Challenging the theoretical framework, a recent review by Gibson et al.^29^ highlighted that studies examining biological markers (e.g., lactate, BDNF), TMS, and behavioral performance separately lack consistent empirical evidence, emphasizing the need for integrative research to evaluate these markers within a unified protocol.

The aim of the present study was to examine (I) the changes in motor performance following explicit SML acquisition through PP or MI, (II) their neurophysiological correlates, (III) and the impact of HIIE on early- and late-consolidation phases (after HIIE, +24 hours and +7 days), along with their underlying biological and neurophysiological substrates. For this purpose, participants were randomly assigned to one of 4 groups: PP_HIIE_, PP_REST_, MI_HIIE_ or MI_REST_, differentiated by the type of practice (i.e. PP or MI) followed by either a rest period or a HIIE. Motor learning was assessed behaviorally, with neurophysiological and biological correlates examined via TMS and blood sample analyses (see Fig. 1).

**Figure 1.**
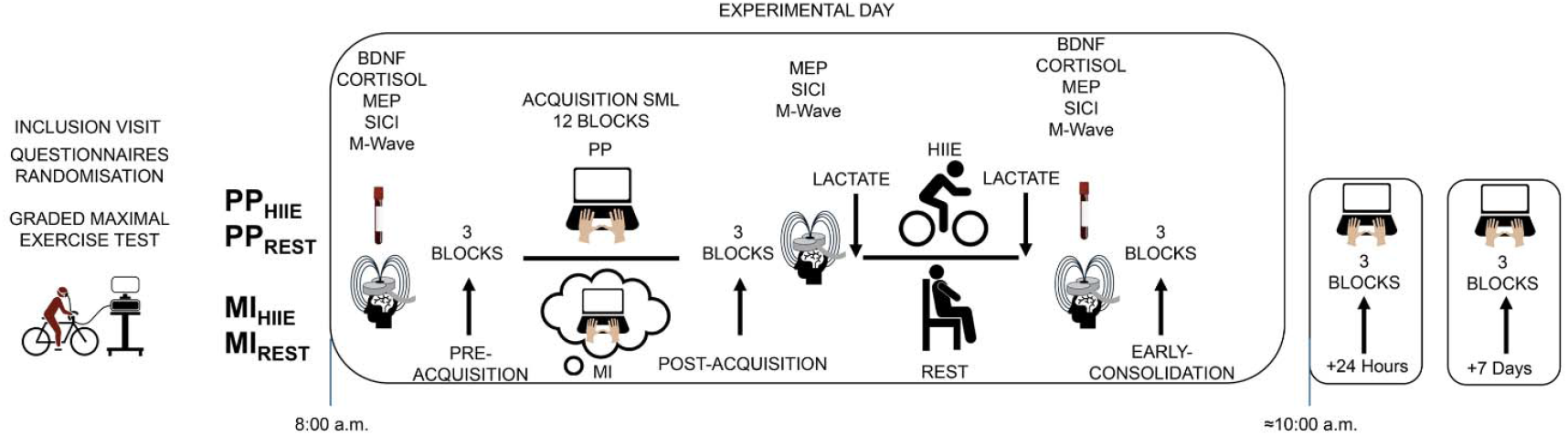
Overview of the experimental design. SML performance was assessed before and after PP and MI acquisition (PRE- and POST-ACQUISITION), and after the HIIE or REST (EARLY-CONSOLIDATION). Behavioral consolidations were tested 24 hours and one week later (+7 days). Blood samples for BDNF and lactate were collected at PRE-ACQUISITION and EARLY-CONSOLIDATION, and MEPs and SICI were assessed at PRE- and POST-ACQUISITION, and EARLY-CONSOLIDATION.

## Results

### Physical and mental practices on SML performance during acquisition

MOVEMENT TIME demonstrated a significant interaction between TIME*PRACTICE 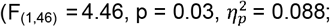 Fig. 2A). Post-hoc revealed a diminution of MOVEMENT TIME for both PP and MI from PRE to POST-ACQUISITION (both p < 0.001). The acquisition-induced decrease in MOVEMENT TIME [%] was significantly greater for PP (46.8 % ± 12.2) compared to MI 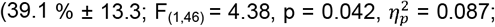Fig. 2B). ACCURACY demonstrated no significant TIME*PRACTICE interaction 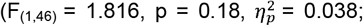 Fig. S1A) nor a main TIME effect 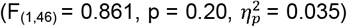, nor a PRACTICE effect 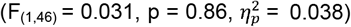 . Acquisition-induced changes in ACCURACY [%] were not different between PP (3.1 % ± 24.4) and MI 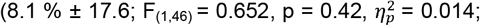Fig. S1B). The learning curve for MOVEMENT TIME is available in the supplementary data (Fig. S2).

### Deleterious impact of HIIE on early- and late SML consolidation

MOVEMENT TIME did not show a main effect of PRACTICE 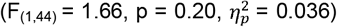 or INTERVENTION 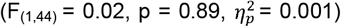, nor an INTERVENTION*TIME 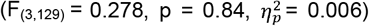, TIME*PRACTICE 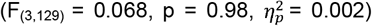, TIME * PRACTICE * INTERVENTION interaction 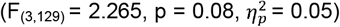. Only a main effect of TIME 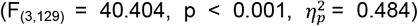 was observed, post-hoc revealing faster MOVEMENT TIME to complete a sequence from POST-ACQUISITION to +24 Hours (p < 0.001), from EARLY-CONSOLIDATION to +24 Hours (p < 0.001), and from +24 Hours to +7 Days (p < 0.001), but no significant difference was observed from POST-ACQUISITION to EARLY-CONSOLIDATION (p = 0.57; Fig. S3). Changes [%] over time were not significantly different between PP and MI or between HIIE and REST (See supplementary data: Fig. S4 B, C, D). While ACCURACY was not influenced by TIME 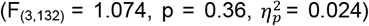 or PRACTICE 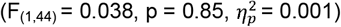, there was a significant main effect of INTERVENTION 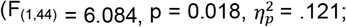 Fig. 2C), indicating that ACCURACY was significantly lower in HIIE groups compared to REST groups. This outcome is further supported by analyses of ACCURACY changes (see below). However, there were no significant interaction TIME*PRACTICE 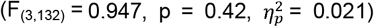, TIME*INTERVENTION 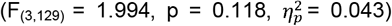, PRACTICE*INTERVENTION 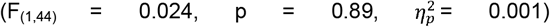 nor TIME*PRACTICE*INTERVENTION 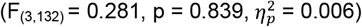 .

**Figure 2.**
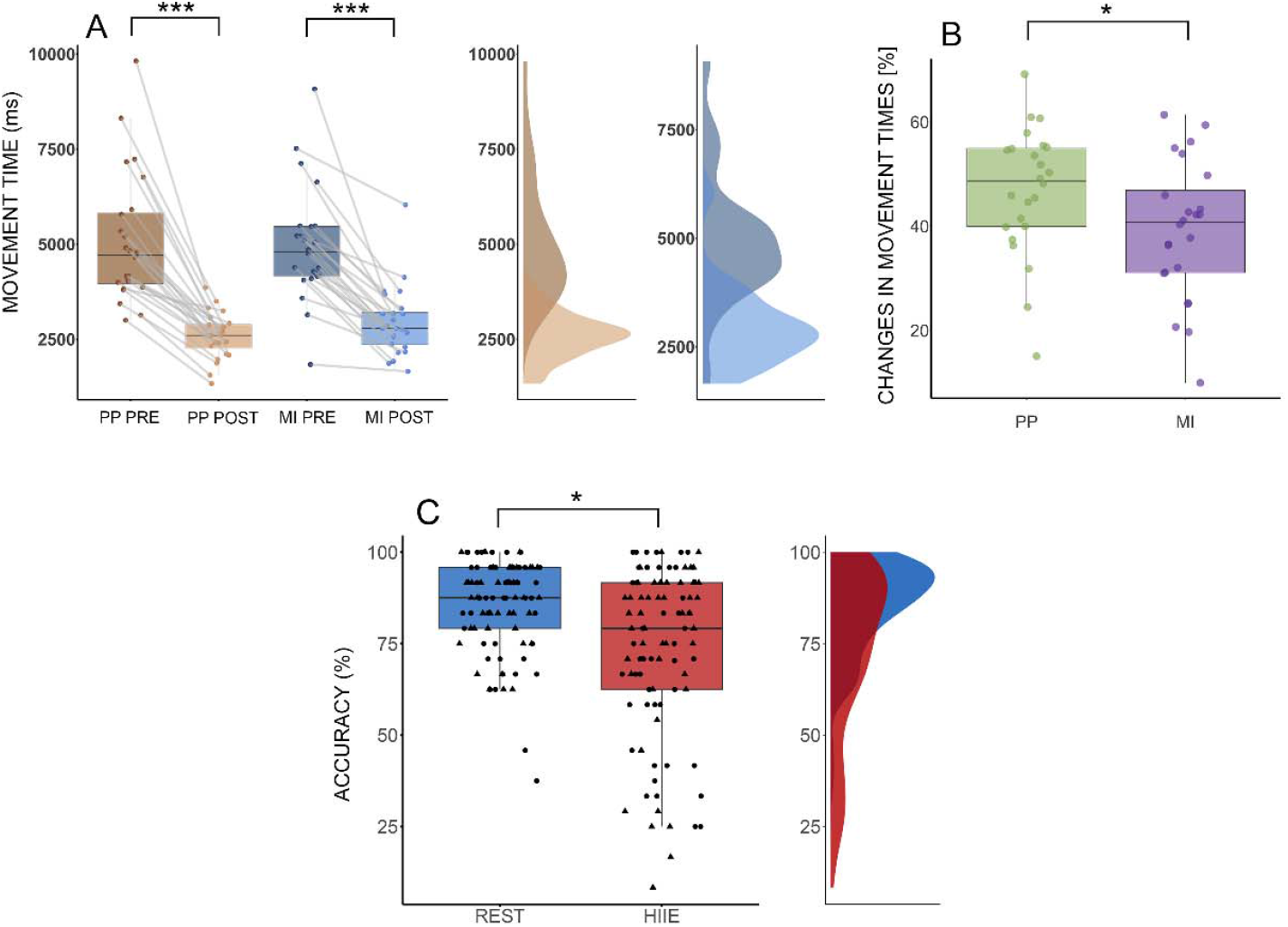
Modulation in MOVEMENT TIME and ACCURACY in SML performance for acquisition and consolidation. A) Performance improvement illustrated by a reduction in MOVEMENT TIME to produce a correct sequence for both PP and MI from pre- to post-acquisition. B) Greater performance gain in MOVEMENT TIME changes [%] during the acquisition stage for PP (46.8 % ± 12.2) compared to MI (39.1 % ± 13.3). C) ACCURACY according to the INTERVENTION factor, including assessment times: post-acquisition, early-consolidation, +24 Hours, and +7 Days. ACCURACY was impaired after HIIE but not REST. Participants in the PP groups are represented through black dot while participants in MI groups are illustrated with triangle. * p < .05; *** p < .001.

Changes [%] in ACCURACY from POST-ACQUISITION to EARLY-CONSOLIDATION revealed a significant impairment in the SML accuracy following HIIE (-12.2 % ± 21.7) compared to REST 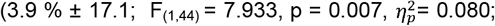Fig. 3B). Similar impairments were observed during later consolidation tests, with a decreasing trend at +24 Hours of consolidation 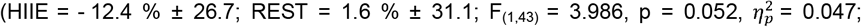Fig. 3C) and a significant decrease at +7 Days 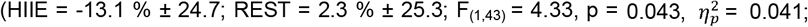 Fig. 3D). It is worth noting that there was no difference in ACCURACY between HIIE (8.1 % ± 25) and REST (3.1 % ± 16.8) from PRE- to POST-ACQUISITION (p = 0.43; Fig. 3A).

**Figure 3.**
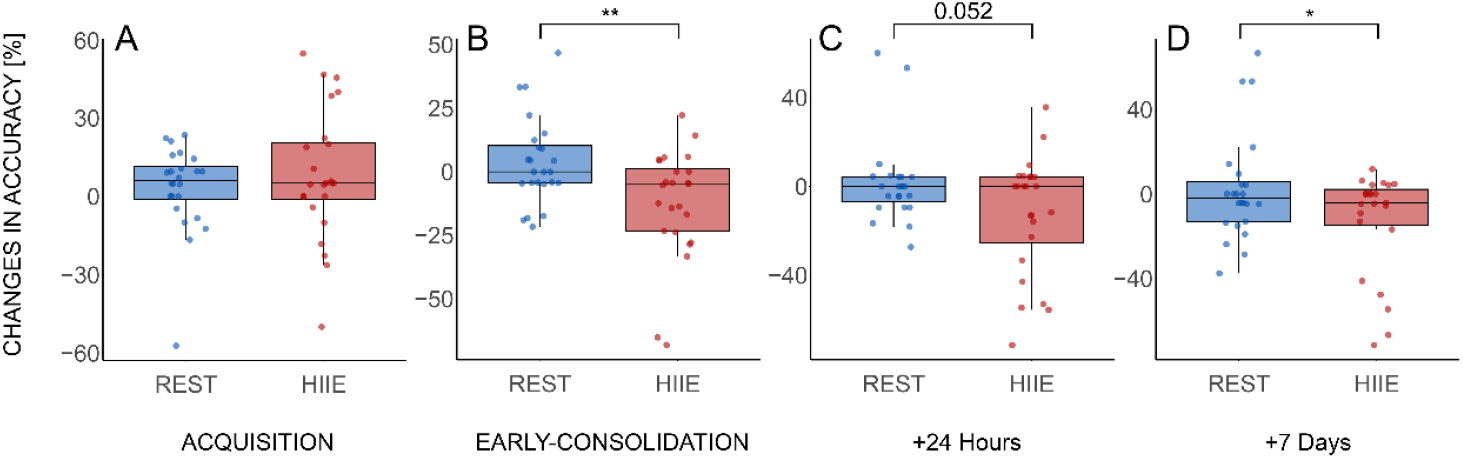
Changes in ACCURACY from the acquisition to early and late consolidation. A) Changes in ACCURACY in % from PRE- to POST-ACQUISITION for REST (i.e. PP_REST_ and MI_REST_ pooled) vs HIIE groups (i.e. PP_HIIE_ and MI_HIIE_ pooled). B, C, and D) Lower ACCURACY at consolidation after HIIE compared to REST. Percent changes from POST-ACQUISITION to EARLY-CONSOLIDATION, +24 Hours, +7 Days tests. * for p < .05; ** for p < .01

### Neurophysiological changes after both PP and MI acquisition, but not after HIIE

Following PP and MI acquisition, MEPs were not significantly influenced by TIME 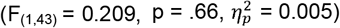, PRACTICE type 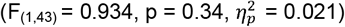, or a TIME*PRACTICE interaction 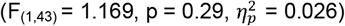. In contrast, SICI showed an effect of TIME 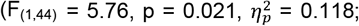 Fig. 4A), regardless of PRACTICE type 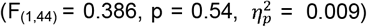 or the TIME*PRACTICE interaction 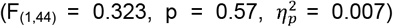. At consolidation, MEPs showed no effect of PRACTICE type 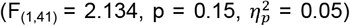, TIME 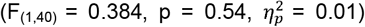, INTERVENTION 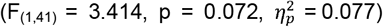, or any interactions (TIME*PRACTICE, 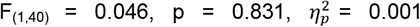,PRACTICE*INTERVENTION, 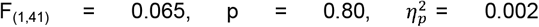, nor TIME*PRACTICE*INTERVENTION,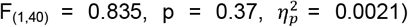. Similarly, SICI revealed no effect of PRACTICE type 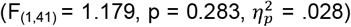, TIME 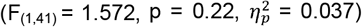, INTERVENTION 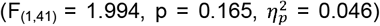, or interactions (TIME*PRATICE, 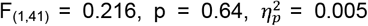 PRACTICE*INTERVENTION, 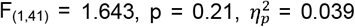 TIME*PRACTICE*INTERVENTION, 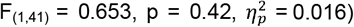.

### Increase in lactate and BDNF levels after HIIE

Absolute changes in lactate concentration from PRE-ACQUISITION to post-HIIE or REST were significantly greater after HIIE compared to REST 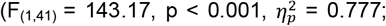 Fig.4B), and a similar pattern of result was found for BDNF concentration 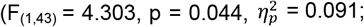Fig. 4C).

**Figure 4.**
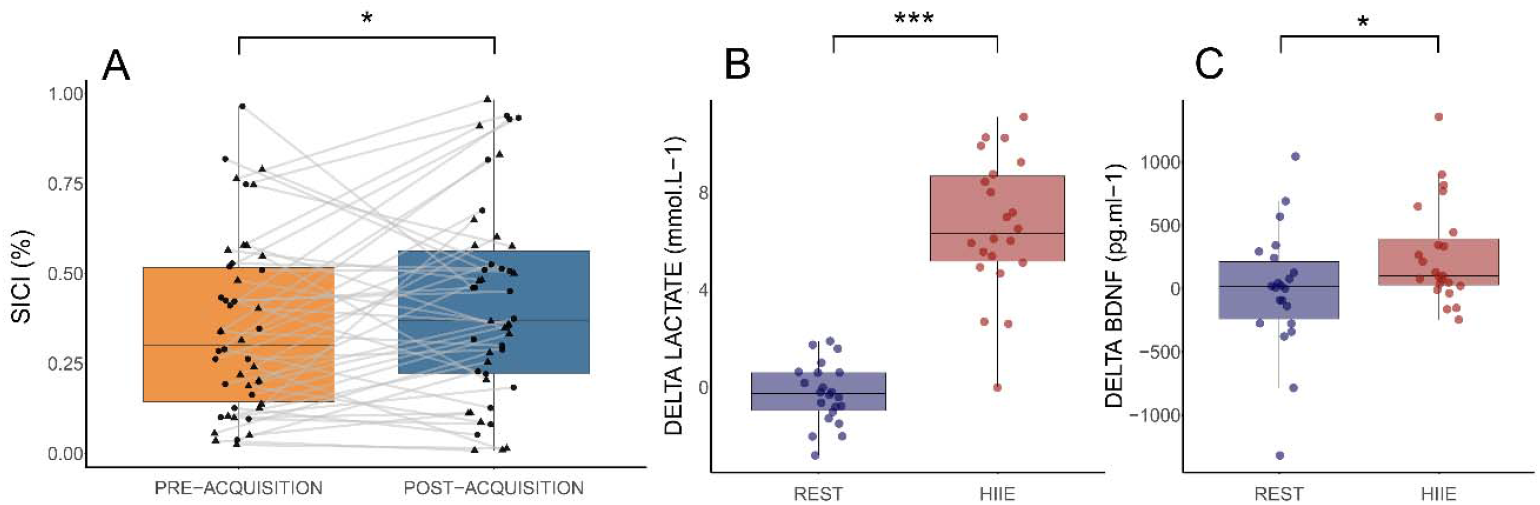
Neurophysiological and biological modulation. A) Both MI and PP elicited SICI increased from PRE-and POST-ACQUISITION of SML (in percent test MEPs amplitude). PP are black dot, MI are black triangle. B and C) Elevation in lactate and BDNF changes for HIIE compared to REST groups, respectively. * p < .05, *** p < .001

### No correlations between behavioral, biological and neurophysiological markers

The analyses of correlations between changes in lactate, BDNF levels, and behavioral performance in ACCURACY during EARLY-CONSOLIDATION showed no significant relationships (respectively p = 0.19, R = 0.2 and p = 0.27, R = 0.1). Changes in MEPs and SICI from POST-ACQUISITION to EARLY-CONSOLIDATION did not correlate with ACCURACY changes at EARLY-CONSOLIDATION (respectively, p = 0.72, R = -0.06 and p = 0.5, R = 0.1). Changes in lactate and BDNF were also not correlated with MEPs changes (respectively p = 0.33, R = -0.15 and p = 0.28, R = -0.17) nor with SICI changes (respectively p = 0.44, R = 0.13 and p = 0.26, R = 0.19). Finally, no significant correlation was found between lactate elevation and BDNF increased (p = 0.21, R = 0.2). All correlations between each time point of behavioral assessments and blood marker levels as well as neurophysiological aspects are presented in the supplementary data (Table S1).

## Discussion

This study aimed to investigate the impact of HIIE on early- and late-consolidation of an explicit SML task acquired with PP or MI, while also examining the associated neurophysiological and biological correlates. We found that both type of practice elicited gains in performance during acquisition, with an advantage of PP that led to greater improvement compared to MI. The acquisition of the SML task with both practices led to a short intracortical disinhibition without changes in corticospinal excitability. The main finding of this study is that motor accuracy in SML was impaired during both early- and late-consolidation following HIIE, regardless of the type of practice. No further changes in corticospinal excitability nor intracortical inhibition were observed after HIIE, despite significant increases in both lactate and BDNF levels.

The first finding of this study is that both types of practice improved movement time, with PP leading to significantly greater improvements in SML acquisition compared to mental practice. These pattern results are commonly described in the field of motor learning with MI^30^. It is well established that PP and MI share common neural substrates and operate within the framework of the internal model theory^31,32^. PP leverages sensorimotor feedbacks to continuously refine and update motor control, whereas MI, in the absence of such feedbacks, induces greater variability and slower adaptation processes, ultimately resulting in reduced performance in motor skill paradigms^30^. At the neurophysiological level, our data revealed that neither PP nor MI modulated MEPs following acquisition, although both decreased SICI. The lack of change in MEPs is not surprising, as literature on motor skill acquisition with PP reports mixed findings, included increased MEPs^7,8^, or decreased MEPs^9,10^, and less commonly no changes^11^. Conversely, the absence of MEPs changes following SML with MI aligns with previous studies reporting similar findings in tasks such as thumb abduction or sequence finger movement opposition^14,33^. In contrast, SICI is considered as a more precise and reliable indicator of the intracortical modulation associated with motor skill acquisition^11^. Here, SICI decreased after both types of practice, a finding consistent with previous reports in the motor learning literature for PP^11,12^, but, to our knowledge, unprecedented in the context of MI practice^15^. The reduction in SICI reflects a modulation in GABAergic activity^34^, a major process supporting motor skill acquisition^35^.

Therefore, while PP yielded greater performance gains than MI, both practices exhibited a consistent pattern of brain excitability following explicit SML acquisition, with unchanged MEPs and decreased SICI, highlighting SICI as a key marker of neuroplasticity independent of practice type.

The major result of this study is that HIIE had a detrimental effect on accuracy at both early and late stages of consolidation regardless of the type of practice. This result contrast with findings from recent studies using modified explicit SML paradigms, which reported no impact of HIIE on motor performance during consolidation^20-22^. The main distinction with motor paradigms that may benefit from HIIE seems to lie in the implicit versus explicit nature of the motor skill practice. Accordingly, it has been suggested that HIIE may disrupt brain networks underpinning the explicit components of SML, particularly the prefrontal cortex which is involved during acquisition and consolidation^20,36^. This explanation is consistent with the Reticular-activating hypofrontality model^37^, which posits that the high resource demands of intense exercise induce hypoactivity in frontal brain regions, thereby impairing the prefrontal cortex involvement to effectively support explicit components of motor skill consolidation. In the studies by Swarbrick et al.^21^ and Cristini et al.^20^ a modified version of explicit SML paradigm with 12-item was used, potentially exceeding the working memory capacity required to explicitly acquire the goal-directed and movement-based components of SML^38,39^. This task ultimately led participants to adopt a mixed strategy, combining implicit and explicit learning which might have reduced susceptibility to consolidation impairments after HIIE. This hypothesis is partially supported by the fact that a large number of participants in Cristini’s study^20^ either failed to recall the sequence in the explicit groups or, discovered part of the sequence in the implicit group. Conversely, Frisch et al.^22^ used a SML task with only 5-item paired with concurrent word list learning, a procedural and declarative learning overlap that is well-documented to interfere when performed in close temporal proximity, potentially explaining the absence of HIIE effects on SML consolidation^40^. Here, we hypothesize that the explicit bimanual 8-item SML task, which involves the prefrontal cortex during acquisition and consolidation tests^41^, may have been particularly susceptible to HIIE-induced disruption of brain networks supporting goal-directed components of SML, ultimately leading to greater performance deterioration after HIIE. Previous studies focused on PP; however, preliminary data in the MI domain, using an explicit unimanual 6-item SML task learned with MI and followed by moderate-intensity exercise, demonstrated benefits in motor performance at +24 Hours of consolidation^23^. This explicit unimanual 6-item task, being simpler and more readily automated than the complex bimanual 8-item task, may have facilitated a faster transition to implicit processing, thereby benefiting from moderate-intensity exercise, which may have also reduced its impact on prefrontal activity. These results from the study by Monany et al. ^23^ should be interpreted with caution, as the small group of subjects showing improved performance at consolidation was the lowest-performing group initially, with the greatest potential for improvement. Altogether, HIIE does not facilitate SML consolidation, irrespective of whether the task is acquired through PP or MI, and its detrimental effects seem to be influenced by the task’s explicit complexity, as observed with the bimanual 8-item task.

Finally, we observed an increase in peripheral blood concentrations of lactate and BDNF following HIIE, a finding consistent with the existing literature^27^, but we did not find any changes in MEPs or SICI after HIIE. This lack of change in MEPs and SICI after HIIE is consistent with the review by Turco and Nelson^42^ which highlights mixed results for these two neurophysiological components following HIIE. Some authors have attributed the beneficial effects of physical exercise to the release of exerkines, such as lactate and BDNF, which promote an environment conducive to brain plasticity and motor learning^43^. While one study has reported a correlation between lactate and BDNF release following an HIIE protocol and improved motor performance during the consolidation phase^27^, other studies have failed to replicate such results^44,45^, as in here. Our findings highlight the ongoing challenge in establishing a clear relationship between biological and neurophysiological modulations following HIIE and motor skill learning. This issue was recently reviewed by Gibson et al.^29^, who noted considerable methodological heterogeneity across study and results within the limited existing literature, and concluded that the role of lactate in modulating corticospinal excitability remains unclear and that other factors may also be involved, thereby, emphasizing the need for further research.

Our study provides evidence that while PP is more effective than MI in enhancing motor performance during acquisition, both practices induce markers of learning-induced neuroplasticity, evidenced by short intracortical disinhibition. The major finding of this study is that practicing HIIE immediately after acquiring an explicit bimanual SML task with physical or mental practice, has a deleterious impact on motor consolidation. This result represents an important step toward understanding the effects of HIIE on motor learning, delineating its positive impact as limited by the complexity of explicit components of the SML to be consolidated. Finally, the mechanistic model of exercise-induced brain plasticity through biological released factors, supposed to trigger changes at the behavioral level, should be reconsidered as the empirical examinations performed here do not support this framework, at least within the domain of explicit SML.

## Materials and Methods

### Ethics information

The present research complies with the ethical regulations, has been approved by the ethical committee of Île de France X (2023-A00155-40), and was conducted in accordance with the Declaration of Helsinki. The study was preregistered on clinical trial (NCT05910814). Once the procedure was fully explained, participants gave written informed consent before inclusion. All participants received an attendance fee. The sample size was determined using G*Power software (version 3.1.9.7) based on an a priori power analysis aimed at detecting an effect (Standardized mean difference) of 0.62, as reported in the meta-analysis by Wanner et al.^18^ (2020), which examined the effect of physical exercise performed immediately after the acquisition phase. The statistical power and alpha level were set at 0.9 and 0.05, respectively, yielding an estimated sample size of 44 participants.

### Participants

Forty-eight right-handed healthy subjects were enrolled for this study and randomized into four groups (PP_HIIE_, PP_REST_, MI_HIIE_, MI_REST_) characterized by the type of practice (PP or MI) and the type of intervention (REST or HIIE). To be eligible, participants had to be aged between 18-35 years old, determined as right-handed according to the Edinburg Handedness Inventory (cut-off > 0.4 for right-handedness^46^), and classified as good sleepers as ensured by the Pittsburgh Sleep Quality Index (PSQI, cut-off < 10;^47^); participants characteristics are displayed table 1. Exclusion criteria included: the presence of ferromagnetic metallic implant, previous neurosurgery, history of seizures, major head trauma, cardiovascular and/or respiratory disease, alcoholism, drug addiction, history of orthopedic problems (upper-lower limbs < 6 months), and any other psychiatric, neurological disorders, or a resting heart rate > 100. Musicians and professional typists were not included to avoid testing participants with a high level of finger dexterity. Participants with an individual scoring < 4 on the Corsi test used to assess the visuo-spatial memory span were also excluded.

**Table 1.**
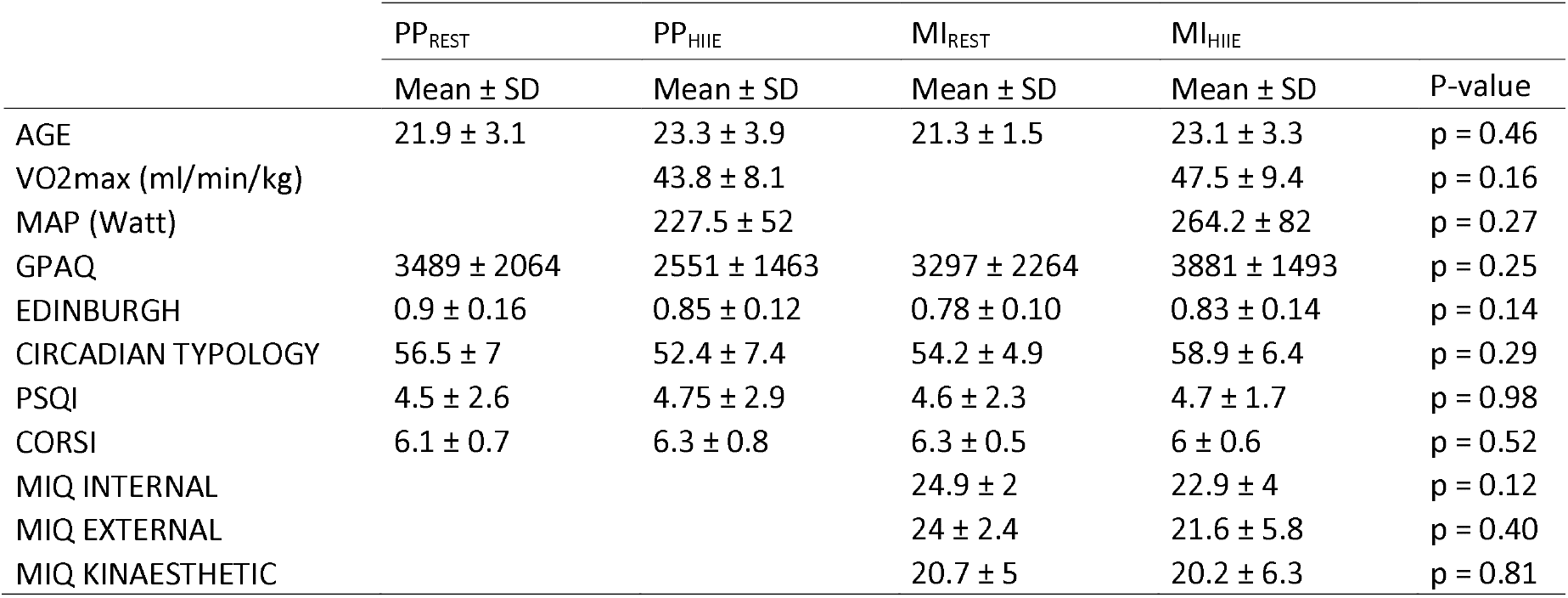
Characteristics of the participants for each group. P-values correspond to the results of Kruskal-wallis or Wilcoxon tests. GPAQ = global physical activity questionnaire; MAP = maximal aerobic power; MIQ = motor imagery questionnaire testing internal, external and kinaesthetic modalities; PSQI = Pittsburgh Sleep Questionnaire Index.

### The bimanual sequential finger tapping task

Participants were seated on a chair positioned 50 cm in front of a computer screen. The fingers were placed on the alphabetical AZERTY keyboard, with Q corresponding to the left little finger, Z to the left ring finger, E to the left middle finger, R to the left index, U to the right index, I to the right middle finger, O to the right ring finger, M corresponding to the right little finger and finally, both thumbs were placed on the space bar. The sequence of eight movements required pressing the keys in the following order R-O-E-M-I-Z-U-Q, and systematically validating the sequence by pressing the space bar with the right thumb (i.e. the left thumb was not used). Participants were instructed not to correct conscious errors but instead to press the space bar to start a new sequence. Each experimental block required participants to repeat the sequence eight times as accurately and as quickly as possible. Each block began with the message “Are you ready?” on the screen, followed by a 3-s countdown before the first sequence, and finished with a 20-s rest period. The sequence remained displayed on the screen during blocks and rest periods. The task was performed using Psytoolkit^48^. The order and the timing of each keypress was recorded and used to calculate the ACCURACY (i.e. the percentage of correct sequences achieved within a block), with a maximum of 8 correct sequences per block, and the MOVEMENT TIME (i.e. average time, in milliseconds, to complete a correct sequence in a block).

### Experimental design

Subjects were randomly assigned to one of the four experimental groups: PP_HIIE_, PP_REST_, MI_HIIE_, or MI_REST_ (Fig. 1). During an initial visit after inclusion and randomization, participants assigned to MI groups completed the Movement Imagery Questionnaire-third version (MIQ-3f;^49^) to assess their visual and kinaesthetic MI capacities, and to familiarize themselves with explicit mental rehearsal of movements. Participants allocated to the HIIE groups performed a graded incremental test on a cyclo-ergometer (Ergoline GmbH, Germany). The incremental ramp protocol involved a 15 or 25 W increase each minute, depending on the subject’s physical condition, until exhaustion was reached, and was used to assess maximal aerobic power and corresponding VO2max (Quark CPET system, Cosmed, Italy). During the experimental day, participants were asked to come at the laboratory at 8:00 a.m., approximately 10 days after the initial visit. The experiment was divided into six phases over a one-week period:

#### PRE-ACQUISITION

Blood samples were collected and TMS was delivered to assess MEPs amplitude and SICI. Participants then began the encoding of the SML by watching a video showing the eight-item sequence to be learned, followed by three consecutive attempts to reproduce the sequence on the keyboard. In case of error, participants rewatched the video before trying again. Upon successful encoding, participants completed three tested blocks of eight sequences performed as accurately and as quickly as possible, with 20-s of rest in-between blocks.

#### TRAINING

Participants performed 12 SML blocks of eight sequences each, with a 20-s rest period between blocks. PP groups physically performed the SML task, while MI groups imagined themselves performing the SML using a combination of visual and kinaesthetic imagery modalities. Importantly, MI participants were instructed to avoid any overt finger movements during mental practice, except that of the right-thumb to end each imagined sequence.

#### POST-ACQUISITION

After training, SML performance was assessed over three blocks, such as in PRE-ACQUISITION, followed by MEPs amplitude and SICI assessments.

#### HIIE OR REST INTERVENTION

Participants in the HIIE groups performed a high-intensity exercise on a cycle ergometer (Ergoline GmbH, Germany). The exercise session began with a 2-min warm-up at 50 W, followed by three periods of 3-min of exercise at 80 % of maximal aerobic power, with 2-min active rest periods at 40 % of maximal aerobic power in between. The total duration of the exercise session, including the warm-up, was 17 min. Participants in the REST groups remained seated while watching a documentary for the same duration.

#### EARLY-CONSOLIDATION

Two and three minutes after the HIIE or REST intervention, lactate was collected, while blood samples were collected at 5 min post intervention. MEPs and SICI measurements were then performed. To investigate the direct effect of HIIE or REST on early consolidation, motor performance was assessed during another three blocks of measurements as previously described.

#### LATE-CONSOLIDATION

The following day (+24 HOURS) and after a week (+7 DAYS), at 10:00 a.m. (± 2:00 h), motor performance on three blocks of the SML was assessed remotely from participants’ homes, with instruction to ensure a quiet environment free of distractions.

### Instruments

#### Neurophysiological assessment

Electromyography (EMG) signals were used to record TMS-evoked potentials and to ensure that there was no muscular activation during acquisition through MI. Participants were equipped with surface EMG electrodes (Kendall™ Medi-Trace® solution Ag/AgCl) placed on the right first dorsal interosseous (FDI) muscle. A bipolar assembly was used with the electrodes positioned on the muscle belly and on the proximal phalanx of the right index, and with a reference electrode placed on the right ulnar styloid process. EMG signals were acquired and amplified by a bio-amplifier (ML138, ADInstruments) and converted from analog to digital at a sampling rate of 2 kHz using a PowerLab (16/30-ML880/P, ADIstruments, Bella Vista, Australia). Online filtering of the EMG signals was performed with a bandpass between 20 and 500 Hz, and the signals were analyzed on Labchart 8 software (ADInstruments). To account for potential changes in sarcolemmal excitability when interpreting MEPs amplitude, responses were normalized to the maximal M-wave (Mmax). The assessment of Mmax on the FDI was achieved with a single constant-current stimulation (DS7R; Digitimer, Welwyn Garden City, UK) applied on the right ulnar nerve via a 30-mm anode-cathode bipolar felt pad (Bipolar Felt pad Stimulating Electrode Part number E.SB020/4mm, Digitimer). Stimulation consisted of a single rectangular stimulation pulse of 1-ms duration, with a maximum voltage output of 400 V. Electrical stimulations were first evoked at 5 mA and then increased by 2-mA steps until the Mmax peak to peak amplitude was obtained. MEPs and SICI were assessed using simple or paired pulse TMS over the left M1, respectively. A figure-of-eight coil with 70-mm wing diameters connected to a Magstim 200 magnetic stimulator (Magstim, Whitland, Dyfed, UK) was used. The coil was positioned at a point that is backward and laterally at a 45° angle to the sagittal plane inducing a postero-anterior current in the brain. The position of the coil was adjusted to ensure optimal location for eliciting the greatest MEP amplitude at a given intensity (i.e. 50% of maximal stimulator output). This stance was drawn with a marker on a cap fitted on the participant’s head to provide a reference landmark for each TMS measurement session. The individual resting motor threshold (rMT) was defined as the lowest intensity that evokes three discernible MEPs out of five evoked responses. MEPs were elicited by a single magnetic pulse delivered at 120 % of the rMT. SICI were evoked using paired stimulations, where a conditioning stimulation at 70 % of rMT is followed by a test stimulation at 120 % of rMT, with a 3-ms interval^13^. During each TMS session, 10 MEPs and 10 SICI responses were recorded. The mean peak to peak amplitude of the 10 MEPs and 10 SICI responses for each time of measurement (PRE-ACQUISITION, POST-ACQUISITION, EARLY-CONSOLIDATION) was used for analysis. MEPs amplitude was expressed in percentage to M-max. SICI was quantified as the mean ratio between conditioned responses amplitude and MEPs amplitude.

#### Biological assessment

Blood samples (5 ml) were collected from the antecubital vein and were centrifuged at 3000 x g for 10 min after 1 h of clotting. The plasma was separated from the serum and was kept at – 80 °C. BDNF was analyzed by means of enzyme-linked immunosorbent assay method (ELISA Kit, MyBioSource, San Diego, U.S.A) according to manufacturer instruction. Blood lactate was collected from capillary blood on the finger and data was immediately analyzed using a lactate analyzer device (Lactate Pro 2, Arkay, Kyoto, Japan).

### Statistical analysis

All the following analyses were performed on the R software with the packages {lme4}, {emmean}, {r2glmm}, and {lmer}. For all analyses, outliers > 3*SD from the residuals of the model were removed from the analysis^27^. All significant factors or interactions were tested with Tukey post-hoc. To test for the changes in SML performance and corticospinal excitability during acquisition stage, a linear mixed model (LMM) was first performed with the fixed effects TIME (PRE-ACQUISITION vs POST-ACQUISITION) and PRACTICE (PP vs MI) and subject-specific random effects. As the residuals of the model did not follow a normal distribution, MOVEMENT TIME was transformed with a logarithmic function while ACCURACY, MEPs, and SICI were transformed using an angular transformation. A linear model (LM) was further used to compare PRE- to POST-ACQUISITION changes using this formula [((POST-ACQUSITION – PRE-ACQUISITION) / PRE-ACQUSITION) × 100] in MOVEMENT TIME and ACCURACY between PP and MI.

To analyze modulations in SML performance during consolidation, a LMM was applied to MOVEMENT TIME and ACCURACY (transformed data), with the fixed effects TIME (POST-ACQUISITION, EARLY-CONSOLIDATION, +24 Hours, +7 Days), PRACTICE (PP vs MI), and INTERVENTION (REST vs HIIE) and subject-specific random effects. Neurophysiological measures were analyzed using the same three factors, with TIME limited to POST-ACQUISITION and EARLY-CONSOLIDATION. Absolute changes in lactate (mmol.L-1; mean of both sample collections after REST vs HIIE) and BDNF (pg.mL-1) concentration from PRE-ACQUISITION to EARLY-CONSOLIDATION were compared between REST or HIIE interventions using a LM. As for the acquisition, a LM was further used to compare changes from POST-ACQUISITION to EARLY-CONSOLIDATION, from POST-ACQUISITION to +24 Hours and from POST-ACQUISITION to +7 Days changes in the MOVEMENT TIME and ACCURACY between PP or MI, and REST or HIIE. Finally, exploratory correlations were performed between biological (lactate, BDNF), behavioral (MOVEMENT TIME, ACCURACY), and neurophysiological outputs (MEPs, SICI) at EARLY-CONSOLIDATION, +24 Hours and +7 Days.

## Acknowledgments

The authors sincerely thank Dr. M. PRUDENT, Dr. G. DEGLICOURT, Dr. L. BERTOLINO and Dr. G. CALLIES, as well as the entire department of functional respiratory testing of the Croix-Rousse hospital, for their invaluable support during participant inclusion and their assistance in handling blood samples. We acknowledge the financial support of this research from both the Institut Universitaire de France (IUF), awarded to Ursula DEBARNOT, and the Hospice Civils de Lyon (HCL).

## SUPPLEMENTAL INFORMATION

### Supplementary method

#### Complementary assessments: sleep, MI and fatigue

Questionnaires and visual scales were used during this experiment to control participants’ sleep quality, fatigue, as well as MI ability and quality, we controlled their sleep during the two nights prior to the experimental session using an actigraphy watch (wGT3X-BT, Pensacola, USA). During the inclusion visit, participants completed the Movement Imagery Questionnaire-third version (MIQ-3f, Robin et al., 2020) to assess their visual and kinesthetics MI capacities and to familiarize themselves with the mental rehearsal of movements. On the experimental day, participants of the MI groups were asked across the training session to self-evaluate a Liker-type scale their difficulty (1 = very difficult; 5 = very easy) and quality of imagery (1 = no image/no sensation; 5 = image as clear as seeing/sensation as intense as during actual performance) at the end of the 4^th^, 8^th^ and 12^th^ blocks. To control for the state of wakefulness, the Stanford sleepiness scale (SSS, Maclean et al., 1992) was used before each assessment of motor performance on the SML task (i.e. PRE-ACQUISITION, POST-ACQUISITION, EARLY-CONSOLIDATION, +24 Hours, +7 Days). At the end of the intervention (HIIE or REST), the subjective perceptions of both physical and cognitive workload were assessed via the NASA-TLX questionnaire. Physical exhaustion was further assessed with the rate of fatigue scale (ROF, Brownstein et al., 2021).

### Supplementary results

**Figure S1.**
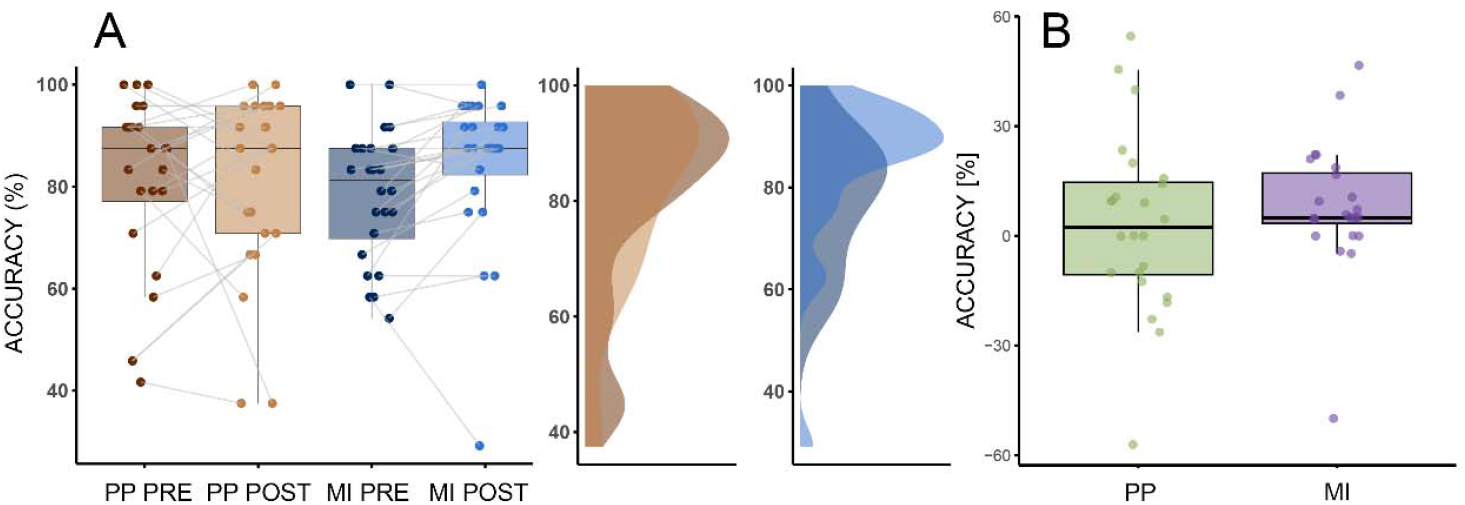
A) ACCURACY according to the type of PRACTICE and TIME of tests. B) Change in ACCURACY [%] from PRE-ACQUISITION to POST-ACQUISITION in both PP and MI.

**Figure S2.**
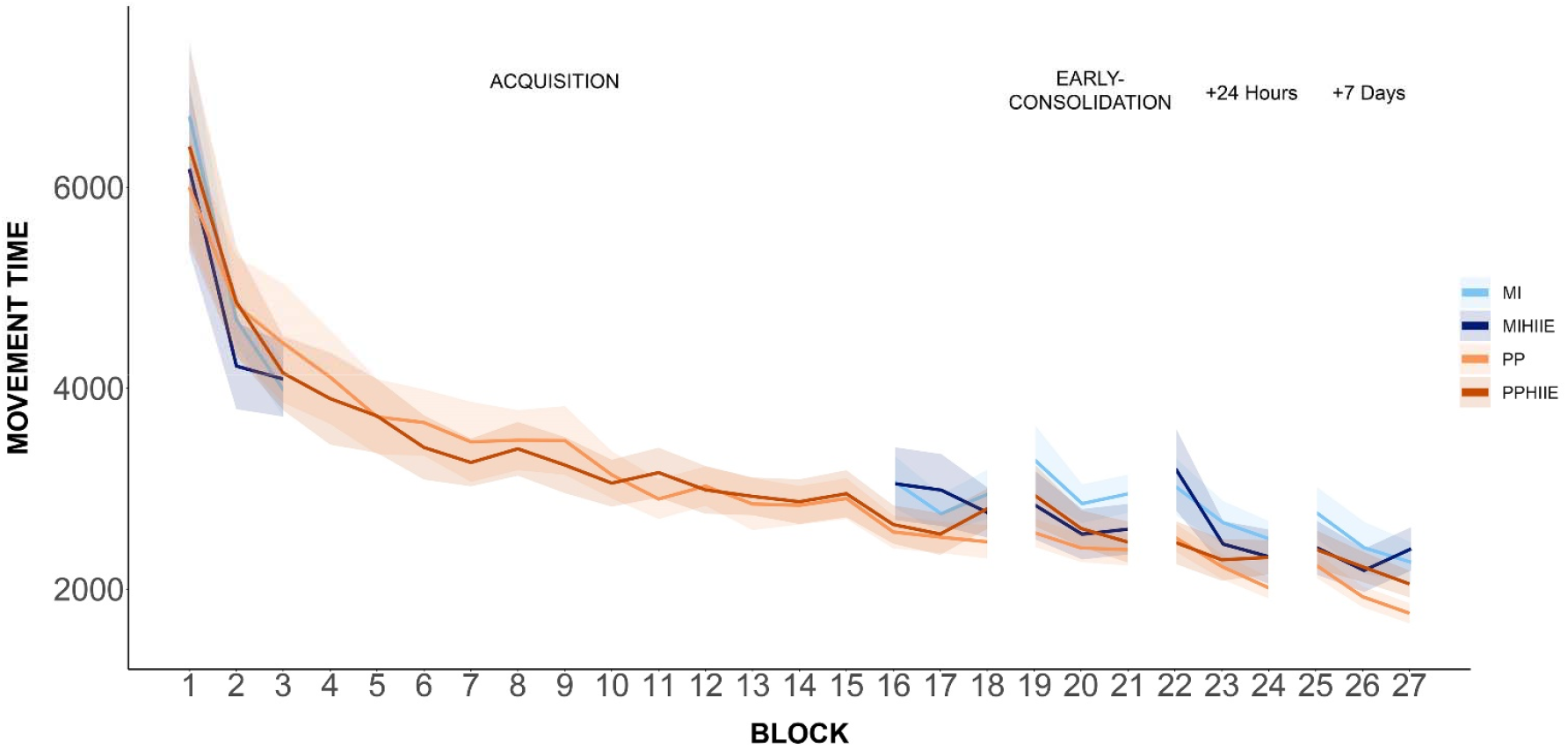
Performance curve for MOVEMENT TIME (ms). Shadow represents the SEM.

**Figure S3.**
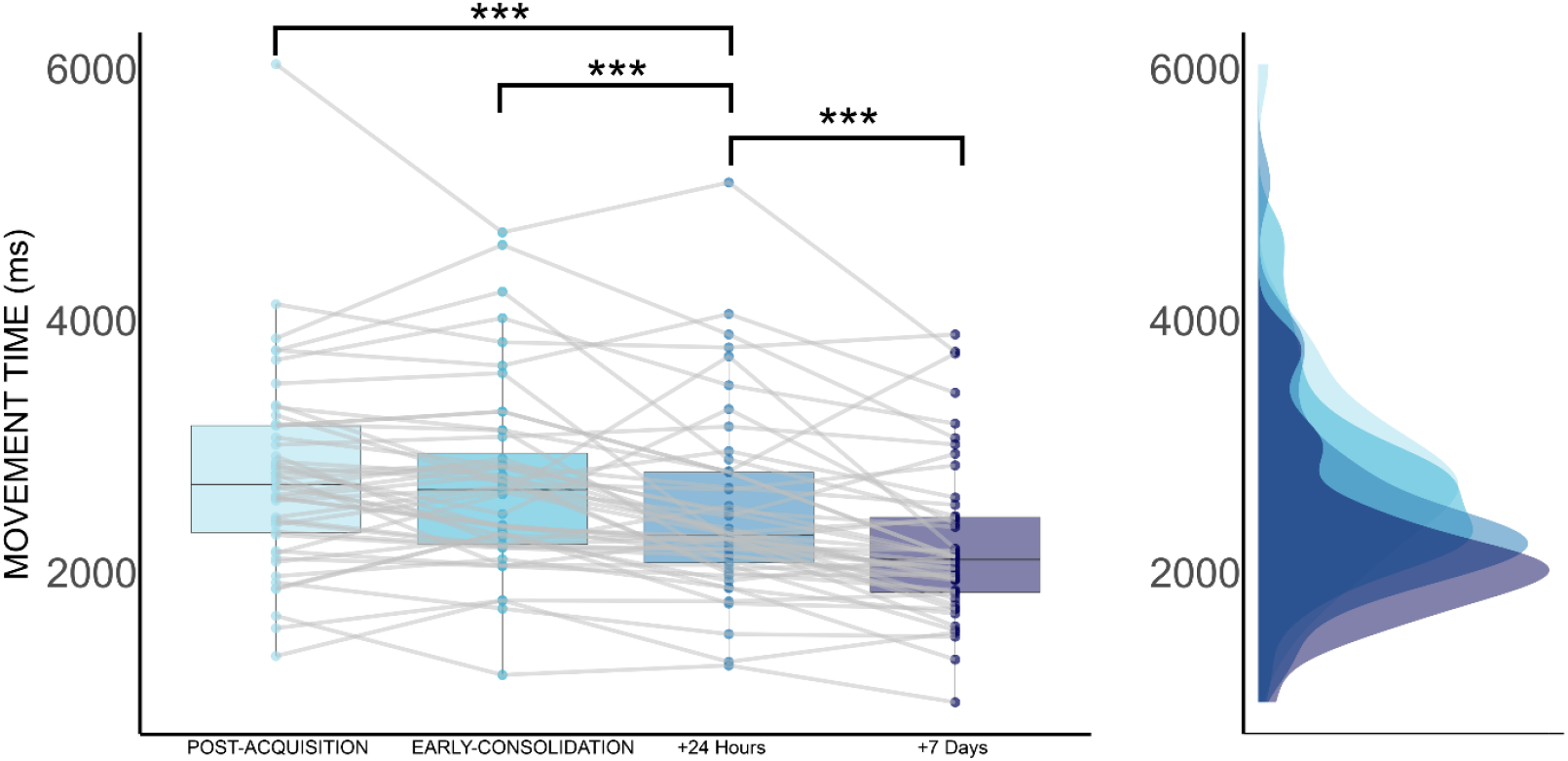
Evolution of MOVEMENT TIME across post-acquisition and consolidation tests. *** p < .001. Changes in MOVEMENT TIME [%] during the EARLY-CONSOLIDATION demonstrated no main effect in the type of PRACTICE (p = 0.49), INTERVENTION (p = 0.24), or an interaction (p = 0.08). For the assessments at +24 Hours, there was no significant main effect of the type of PRACTICE (p = 0.60) and INTERVENTION (p = 0.76) nor the interaction between the factors type of PRATICE and INTERVENTION (p = 0.48). Finally, for the consolidation test at +7 Days, there were no significant main effect of type of PRACTICE (p = 0.75) and INTERVENTION (p = 0.78) nor the interaction between the factors type of PRATICE and INTERVENTION (p = 0.10).

**Figure S4.**
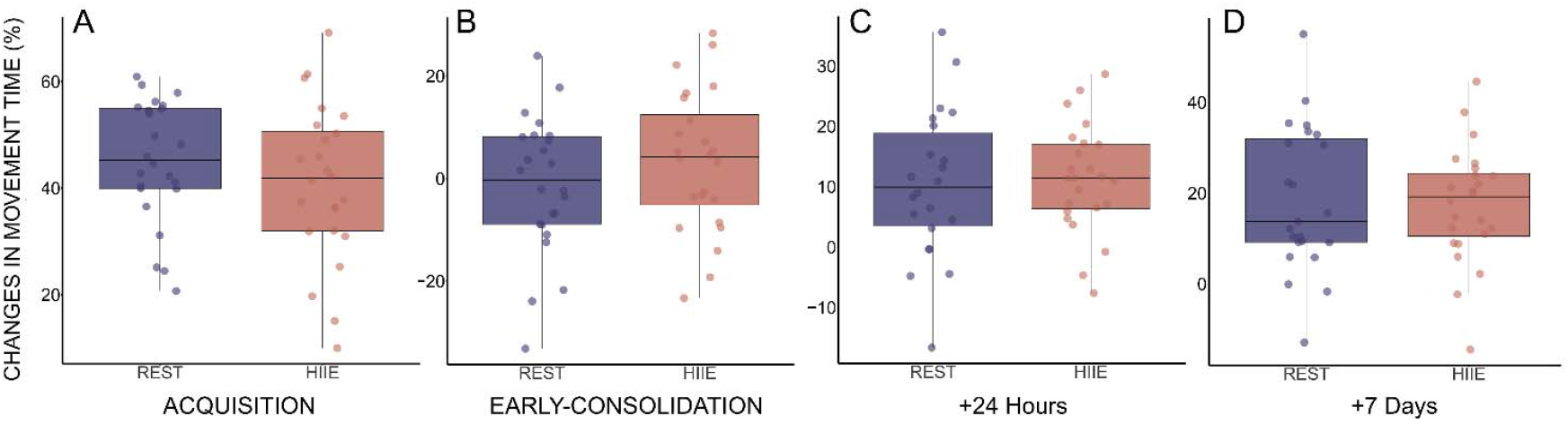
A) Corresponds to the changes in MOVEMENT TIME from PRE-to POST-ACQUISITION for REST vs HIIE groups. Figure S1B, C, and D illustrate the changes [%] in MOVEMENT TIME in relation to the factor INTERVENTION (i.e., HIIE, REST) for each consolidation time point (i.e., EARLY-CONSOLIDATION, +24 Hours, +7 Days), using POST-ACQUISITION as the reference time point.

**Table S1.** Descriptive r and p-value for all correlations between behavioral (changes in percentage), neurophysiological (changes in percentage) and biological analysis (absolute changes).

|  | MOVEMENT TIME |  |  | ACCURACY |  |  | BLOOD |  |  |
| --- | --- | --- | --- | --- | --- | --- | --- | --- | --- |
|  | EARLY-CONSOLIDATION | +24 Hours | +7 Days | EARLY-CONSOLIDATION | +24 Hours | +7 Days | BDNF | LACTATE | CORTISOL |
| MEP | $p = 0.72 / r = -0.06$ | $p = 0.95 / r = 0.01$ | $p = 0.06 / r = 0.29$ | $p = 0.09 / r = 0.26$ | $p = 0.18 / r = 0.21$ | $p = 0.07 / r = 0.28$ | $p = 0.28 / r = -0.17$ | $p = 0.33 / r = -0.15$ | $p = 0.54 / r = 0.1$ |
| SICI | $p = 0.5 / r = 0.1$ | $p = 0.7 / r = 0.08$ | $p = 0.66 / r = 0.12$ | $p = 0.45 / r = -0.12$ | $p = 0.86 / r = 0.03$ | $p = 0.51 / r = 0.1$ | $p = 0.26 / r = 0.19$ | $p = 0.44 / r = 0.13$ | $p = 0.73 / r = 0.06$ |
| BDNF | $p = 0.27 / r = -0.17$ | $p = 0.43 / r = -0.12$ | $p = 0.41 / r = -0.12$ | $p = 0.46 / r = -0.11$ | $p = 0.82 / r = 0.03$ | $p = 0.33 / r = 0.15$ | / | $p = 0.21 / r = 0.2$ | $p = 0.86 / r = -0.03$ |
| LACTATE | $p = 0.19 / r = 0.2$ | $p = 0.75 / r = 0.05$ | $p = 0.72 / r = -0.05$ | $p = 0.24 / r = -0.18$ | $p = 0.25 / r = -0.17$ | $p = 0.1 / r = -0.25$ | $p = 0.21 / r = 0.2$ | / | $p = 0.52 / r = 0.1$ |
| CORTISOL | $p = 0.35 / r = -0.14$ | $p = 0.14 / r = -0.23$ | $p = 0.09 / r = -0.26$ | $p = 0.68 / r = -0.06$ | $p = 0.52 / r = 0.1$ | $p = 0.75 / r = -0.05$ | $p = 0.86 / r = -0.03$ | $p = 0.49 / r = 0.11$ | / |

